# What do we know about survival of Common cranes? An elementary introduction with Euring databank

**DOI:** 10.1101/2021.09.02.458691

**Authors:** Luis M. Bautista, Juan C. Alonso

**Author notes:** **unpublished preprint** submitted to: PROCEEDINGS of the 9th European Crane Conference (2018, Arjuzanx, France). A. Salvi et al. (editors).

## Abstract

The increase of the western populations of Common cranes (*Grus grus*) in the last five decades highlights the need to estimate survival rates. According to Euring databank (EDB), the oldest Common crane ever known was 27 years old in year 2017. This lifespan was obtained by means of 24,900 recoveries of 2,124 ringed cranes collected between years 1936 and 2017. Nearly all cranes were ringed and observed in the last 30 years, and therefore the elapsed time was not enough to reach the maximum longevity reported for the species in captivity (43 years, Mitchell 1911). Life expectancy was five years on average after the ring was attached. Here we provide some elementary analyses to calculate the annual *apparent* survival rate (ϕ = 0.85) and the annual encounter probability (*p* = 0.45) of Common cranes, as a first step to advance in the knowledge of the species’ population dynamics. The great increase of breeding and wintering crane populations in western Europe in the last decades remains largely unexplained.

---

Even though several crane species are threatened or endangered, the world population of Common cranes (*Grus grus*) is likely beyond 700,000 birds (International Crane Foundation 2018). Its population size in western Europe could have increased since the last quarter of the 20th century (Hansbauer et al. 2014), but the mechanisms leading to the increase are not well known or appear contradictory. For example, the increase in numbers of cranes migrating and overwintering can be explained by methodological improvements for counting cranes and also by the increase in numbers of birdwatchers and wildlife managers counting them since 1975 (Alonso et al. 2016). But a better quality and quantity of censuses at the wintering grounds does not account for the increase in the number of breeding pairs reported in northern and central Europe (e.g., Lehrmann and Mewes 2018; Nielsen 2018; Ojaste et al. 2018; Vegvari 2018). It is interesting to note that the number of breeding cranes in Germany, where many pairs have been monitored in their breeding areas for more than 30 years (Prange 2005, 2014), increased six-fold. Overall, the breeding pairs are growing in north-western Europe but apparently not in north-eastern Europe, where they may have even decreased (Prange 2014). A negative density-dependence effect on breeding success (Alonso et al. 1987) could have started to operate at the most populated crane areas in Europe, as suggested by the higher breeding success in regions with a lower breeding density (Prange 2014).

The positive influences of extra food from agriculture, climate, habitat restoration and protection measures must have played a role in the breeding population increase (Leito et al. 2015; Prange 2014) and induced a northward shift of the wintering range in recent decades (Alonso et al. 2016 and references therein). Even a shift from eastern to western Europe, but this possibility remains unproven and there is little evidence that breeding cranes from eastern Europe become breeders in western Europe (Vegvari 2018). Regardless of how the habitat and climate changes have determined the population increase of Common cranes in western Europe, standard demographic metrics as survival or productivity must be known to understand the biological process underpinning such increase.

It is noteworthy that survival data of Common cranes in the western Palaearctic are scarce. We did not find published, recent and detailed survival data about wild Common cranes in western Europe. The first data on longevity (43 years) and average lifespan (12 years, N=7 cranes) of captive cranes were published more than a century ago (Mitchell 1911) and to our knowledge, none was published afterwards. To solve this lack of survival data of Common cranes, we run a trial analysis as a starting point for future, more detailed studies. The first constraint was finding a database large enough for analysis. Some Common cranes have been ringed since the early decades of last century, but public-accessible databases of ring-recovery data are lacking. The only database that is open for researching and publishing is EURING Data Bank (EDB, du Feu et al. 2016). EURING is an acronym standing for European union for bird ringing, a community of national ringing centres that was created to facilitate large-scale analyses of movements and demography of birds. This community welcomes applications from researchers wishing to analyse the EURING data. In 2018 we applied and received 24,900 recoveries of 2,124 Common cranes ringed between 1936 and 2017. Recoveries were encounters of alive or dead cranes since year 1936. According to EDB code of age, most cranes were ringed as nestlings or chicks, unable to fly freely and still able to be caught by hand (EDB code 1; 92.8% of cranes) or full-grown and able to fly freely (EDB code 2; 4,2% of cranes). A small number of cranes were ringed as full-grown individuals hatched in the breeding season of the ringing calendar year, before 31^st^ December and therefore at an age of less than six months old (EDB code 3; 1,3% of cranes). Minimum longevity was calculated as the difference between the ringing date and the last recovery. Longevity was a minimum guess because there were few recoveries of dead cranes (9.8% of cranes).

Very few cranes were ringed in the first 50 years since 1936 but the number increased to 76±9 every year afterwards (mean ±CI95%, 1989-2015). Apparent maximum longevity of 27 years corresponded to a chick ringed in 1989 in Sweden, though EURING reports 24 years as the maximum age also for a Swedish chick (Fransson et al. 2017). Beyond that disagreement it is worth to highlight that maximum lifespan of wild Common cranes calculated with cranes ringed and encountered in 1989 to 2017 (28 years) cannot be biologically meaningful. As expected by the time elapsed since ringing, apparent maximum longevities are blocked in recent years by the moving-wall of time (Fig. 1b). If we take as a working hypothesis that cranes could live in the wild as much as in captivity (43 years, Mitchell 1911), we should wait at least another 15 years starting to count from year 2017. It is true that wild birds typically show shorter lives than birds living in zoos, but here the point is that the maximum age of the cranes ringed every year is blocked by the moving-wall of time, hence there is a problem regarding the maximum lifespan in each year. Awaiting 15 years is required to completely rule out the effects of the moving-wall of time effect on maximum lifespan, at least for cranes ringed in the nineties.

**Figure 1.**
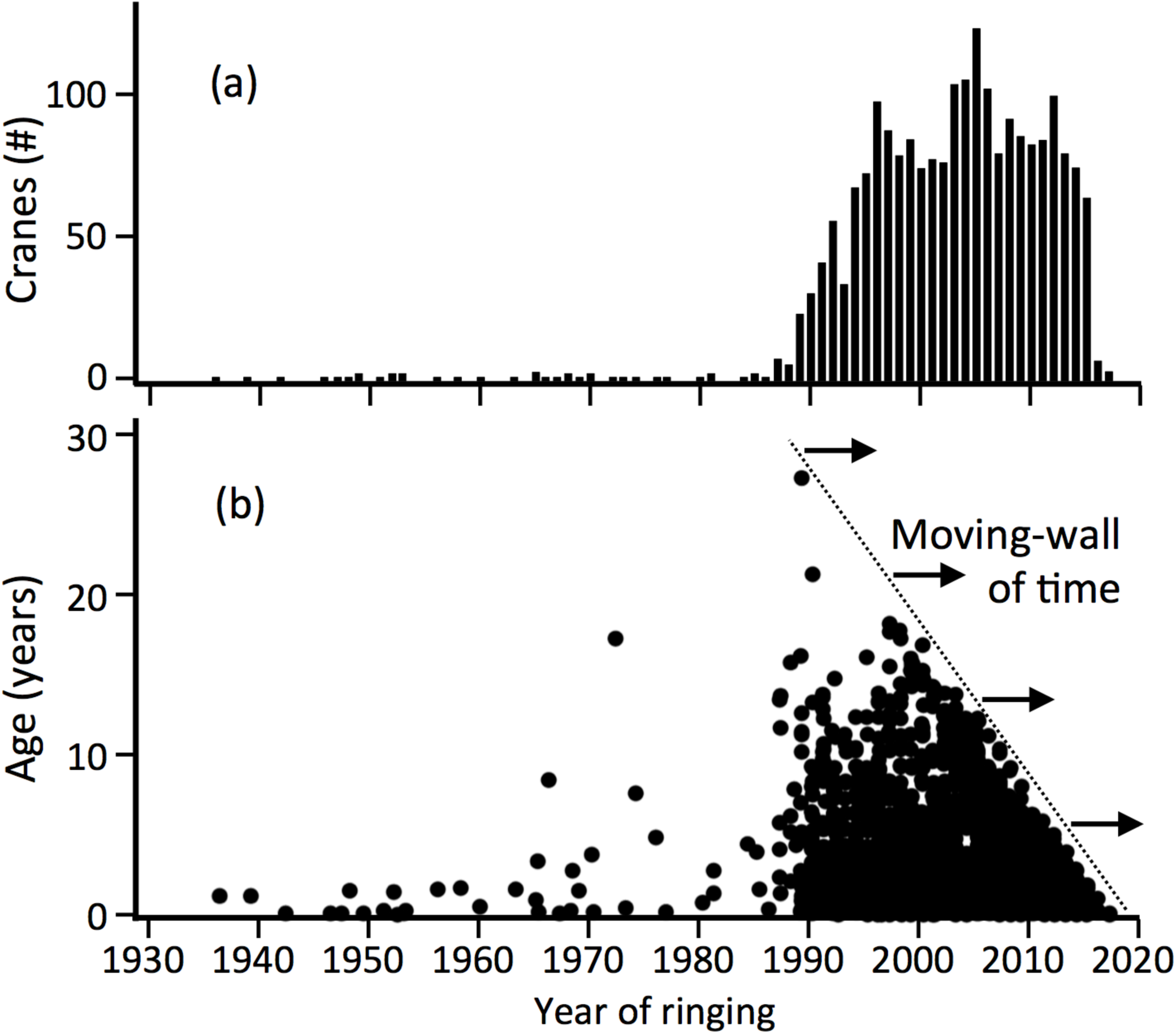
Number of Common cranes ringed each year (a) and apparent age (b) calculated with encounters sent to EURING Data Bank (1936 – 2017). Note that maximum ages in recent years were limited by the elapsed time since they were ringed (moving-wall of time).

Life expectancy in the first year of age (LE_**1**_) calculated as the average number of years remaining at the tagging date must also be biased by the effect of the moving-wall of time, but to a lesser extent than maximum lifespan, because the former is computed with all birds and not just with the long-lived ones. For example, LE_**1**_ did not show a significant temporal trend between 1989 and 2000 (Fig. 2, LE_**1**_ = 4.7±0.8 years, mean±CI95%). The median of LE_**1**_ was 4.1 years (0.3-10.3, IQR 10-90%), a bit smaller than the mean because frequency distribution of ages was asymmetrical. The selected 12-years period is arbitrary, but useful to highlight that some demographic variables of Common cranes can be estimated even if the moving-wall of time is blocking the frequency distribution of lifespan.

**Figure 2.**
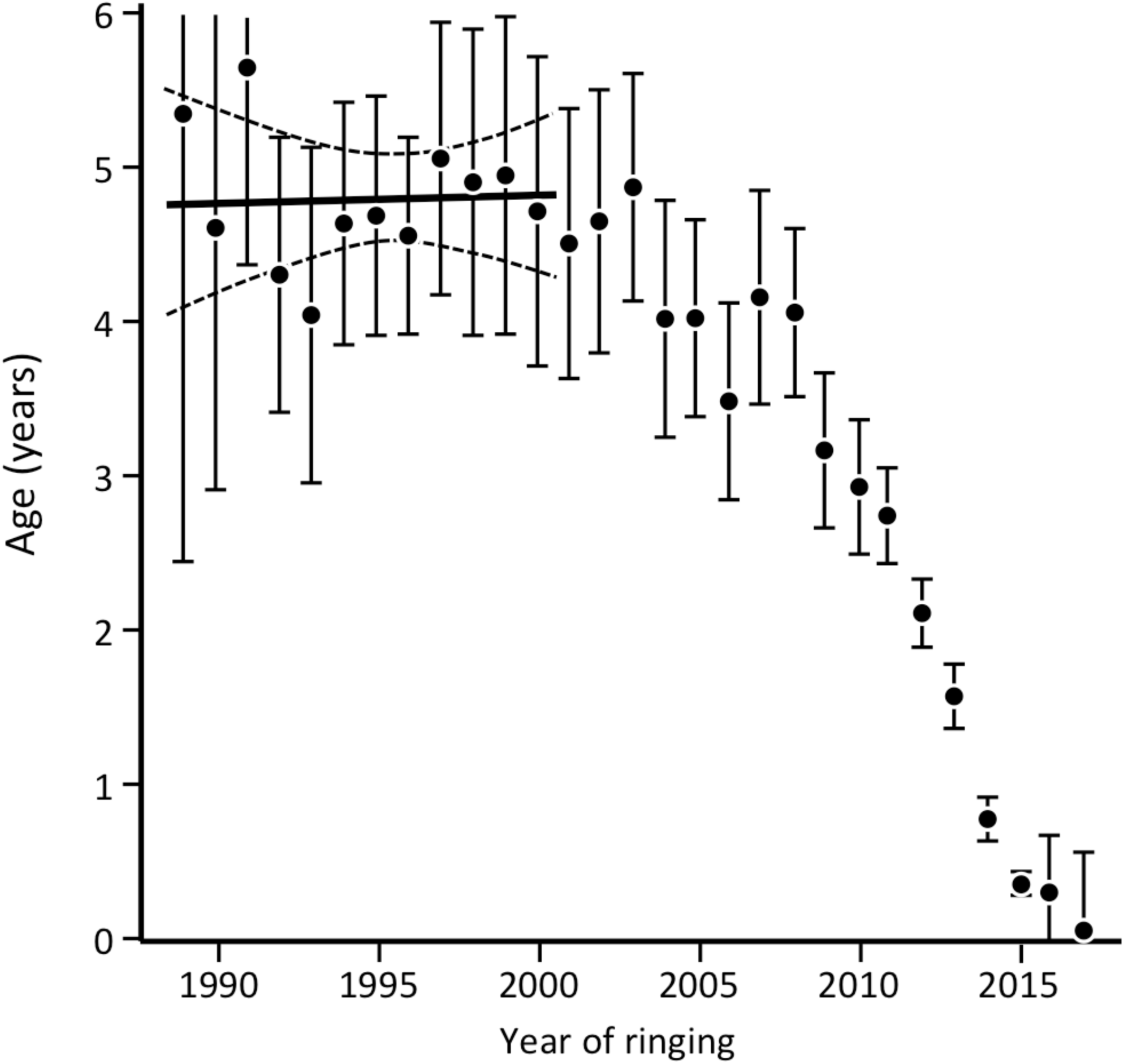
Life expectancy in the first year of life of cranes (Age, mean ± CI95%) by ringing years since 1989. Horizontal thick line and dashed lines show the linear regression (± CI95%) fitted to ages of 741 cranes ringed between 1989 and 2000. The number of cranes in each year is shown in Fig. 1a.

Most cranes included in the calculation of LE_**1**_ survived less than five years because after the last encounter they were not seen again in the following years. But because very few cranes were reported to EDB as found dead, the LE_**1**_ provides just an *apparent* life expectancy (i.e., cranes could be alive but not seen by observers for some time after the last encounter). There are some ways to estimate how long they might be alive after the last encounter. For example, the *long-term* LE_**1**_ could be estimated by the asymptotic curve fitted to the historic increase of the observed mean age for each annual cohort (1989, 1990, etc.). The increase of mean age for each cohort showed a decelerating trajectory as years (*t*) advanced. This trajectory was fitted to each annual cohort from 1989 to 2000 with a mechanistic growth model LE_**1**_(*t*) = θ_1_[1-θ _2_·exp(−θ_3_·*t*)] and solved by the analytic Gauss-Newton method in JMP (SAS Institute Inc. 2015). For example, LE_**1**_ of cranes ringed in 1989 increased the following years according to 5.35[1-0.99·exp(−0.14·*t*)], where all coefficients were statistically significant (p<0.05). Asymptotic LE_**1**_ of cranes ringed in 1989 was 5.35 years (±0.45 CI95%). The mean of all asymptotic LE_**1**_ of cranes ringed between 1989 and 2000 was 4.8 .±0.3 years (mean ±CI95%, N=12 years). This LE_**1**_ ≈ 5 years was similar to that estimated for other crane species as for example Florida Sandhill cranes *G. canadensis pratensis* (mean life expectancy: seven years, Tacha et al. 1992). Although it has been claimed that captive Common cranes can reach a mean age of 12 years (Mitchell 1911), the same mean age that has been calculated for nesting wild cranes through sonographic methods (Wessling 2018, pers. comm.) these mean age values could be the result of exceptional good conditions, either internal or external (i.e., respectively favoured by captivity or territoriality), that may not properly represent the overall population of the species in the wild. We suggest that five years of life expectancy for wild Common cranes is a plausible guess based on the data available in EDB. Every few years the calculations will be updated with more and better data because about 80 new cranes will be added to EDB if the tagging rate holds in future years (Fig. 1a). Besides, the blocking effect of the moving-wall of time over cranes ringed in the first years of the series will decrease.

Most ringed cranes were not encountered in all years, thus the *apparent* annual survival and encounter probabilities must be calculated independently. The encounter probability is useful also to calculate how long we expecte for a live crane to pass unnoticed after its last encounter, a period that for example can be added to the *long-term* LE_**1**_ calculated in the former paragraph. Software MARK (White and Burnham 1999; White and Cooch 2017) provided a likelihood approach to determine the *apparent* survival (ϕ) and encounter (*p*) probabilities. In this elementary introduction we assumed they were both constant across ages and years. Additional studies will introduce age dependence and time dependence, because survival must be smaller in juveniles than in adults and environmental stochasticity predicts variable values for both survival and encounter probability across years (e.g., Nedelman et al. 1987).

The *apparent* annual survival of cranes ringed between 1989 and 2011 (N= 1611 cranes) was ϕ = 0.849±0.008 (±CI95%) and the encounter probability *p* = 0.454±0.013 (±CI95%). This ϕ ≈ 0.85 was comparable to that of similar crane species, as for example the Sandhill crane *G. canadensis* (0.82-0.95, Bennett and Bennett 1990; Fronczak et al. 2015; but see Servanty et al. 2014; Tacha et al. 1992), Whooping crane *G. americana* (0.90, Barzen and Ballinger 2017; Canadian Wildlife Service and US Fish and Wildlife Service 2007) or Red-crowned crane *G. japonensis* (0.80-0.91, Momose 2013). The encounter probability cannot be easily compared to *p* of other cranes species because it is not ordinarily reported, at least up to our knowledge. A *p* ≈ 0.45 means that a ringed crane has a probability of 0.45 of being resighted each year, or in other words, that it will be encountered in about half of the years through its life.

The results highlight the difficulties to explain the population increase of breeding and wintering cranes in western Europe as an intrinsic demographic growth alone. Population growth rates that would result from most published series of crane counts are not possible with the species’ annual recruitment of c. 12% measured in the wintering grounds (Alonso and Alonso 1987; Alonso et al. 1990) and the *apparent* survival rate of ϕ = 0.85 obtained in the present study. Further refinements in calculation of ϕ with EDB or other databases of ringed cranes could rise ϕ, but intrinsic demographic growth alone would be still unlikely even assuming a conservative survival rate of ϕ = 0.92 (see Alonso et al. 2016). Current data on longevity does not contribute to solve the difficulties, because the moving-wall of time blocks the maximum lifespan in EDB records (27 years, Fig.1). Maximum lifespan in the wild must be smaller than in captivity (43 years, Mitchell 1911), but nonetheless accepting 43 years as the top longevity, the life expectancy would be just seven years (ϕ = 0.85), close to the asymptotic LE_**1**_ ≈ 5 calculated in present study with EDB (Fig. 1). A greater survival rate (ϕ = 0.92, Alonso et al. 2016) would increase LE_**1**_ to eleven years, which is close to the life expectancy of 12 years reported in nesting wild cranes by sonagraphical methods (Wessling 2018, pers. comm.). However, maximum lifespan must be notably shorter in the wild than in captivity, hence a value of 43 years seems quite unlike as an estimate of maximum longevity.

In conclusion, some basic questions about demography of Common cranes can be answered with Euring Data Bank. Some parameters, however, like the lifespan in the wild, and other questions, like the recent remarkable population increase observed in western Europe, cannot be solely by the demographic analyses carried out in the present study. The results shown here will be challenged and enhanced in the time coming, because the reports of ringed cranes have increased rapidly in the last decades, through the use of friendly online databases (iCORA, Crane Conservation Germany 2018) and citizen science reports (e.g., Friends of Gallocanta Association AAG 2018).

